# HAT1 Regulates Intestinal Stem Cell Proliferation and Differentiation

**DOI:** 10.64898/2026.03.16.712164

**Authors:** Prabakaran Nagarajan, Caden J. Martin, Andrea R. Keller, Kübra B. Akkaya-Colak, Maria H. Festing, Maria M. Mihaylova, Mark R. Parthun

## Abstract

Stem cells are critical for the development and maintenance of tissue integrity. An important example is intestinal stem cells (ISCs) that generate all epithelial cell types necessary for formation of the intestinal lining. HAT1, a histone acetyltransferase that acetylates newly synthesized histone H4 molecules on lysine residues 5 and 12 during replication-coupled chromatin assembly, is specifically expressed in intestinal stem and progenitor cells located in intestinal crypts. To determine if HAT1 is important for intestinal stem and progenitor cell function, we generated an inducible deletion of the HAT1 gene in intestinal epithelial cells. Loss of HAT1 resulted in morphological defects in the proximal end of the small intestine. Following loss of HAT1, intestinal crypts became elongated, with an increase in stem and progenitor cell proliferation and an increase in the population of OLFM+ cells. Loss of HAT1 also resulted in alterations in intestinal stem cell differentiation, including an increase in the number of Goblet cells and the mislocalization of Paneth cells into villi. HAT1 is specifically responsible for the acetylation of histone H4 lysine 5 (H4K5ac) in intestinal stem cells. Genome-wide characterization of HAT1-dependent H4K5ac in intestinal crypt cells indicates that the most significant loss of H4K5ac occurs in lamina-associated domains (LADs). Loss of H4K5ac in LADs is accompanied by an increase in histone H3 K9 tri-methylation indicating that HAT1 regulates LAD chromatin structure in intestinal crypt cells. A direct role for HAT1 in intestinal stem cell function was demonstrated using organoids in culture. HAT1 is required for differentiation in organoids and for the maintenance of Lgr5+ stem cells. These results indicate that HAT1 is required for the proper regulation of intestinal stem cell renewal and differentiation.

## Introduction

Epithelial cells of the intestine are organized along the crypt-villus axis. At the base of the villus is a structure known as the crypt that is the location of the stem cells and their niche. As the stem cells divide, cells that will differentiate move up the crypt into the transit amplifying zone where they become progenitor cells. The intestinal stem and progenitor cells differentiate into 2 lineages: the secretory and absorptive. The secretory lineage includes Paneth, goblet, tuft and enteroendocrine cells, while the absorptive lineage is comprised primarily of enterocytes. Cells become fully differentiated as they leave the transit amplifying zone and move into the villus. The proper regulation of intestinal stem cell (ISC) proliferation and differentiation is essential to maintain intestinal function^1–3^ under homeostasis as well as rapid responses to injury

ISC proliferation and differentiation is regulated by a complex network of signaling pathways that control cell type specific gene expression patterns^2,4–7^. Unsurprisingly, chromatin structure and epigenetic regulators have been found to play integral roles in the function of ISCs^6,8–10^. Many of these factors play central roles in the repression of transcription. These include components of the polycomb repressor complexes 1 and 2 (PRC1 and PRC2), which are required for facultative heterochromatin, the histone H3 lysine 9 methyltransferase Setdb1, the histone deacetylases HDAC1. HDAC2, HDAC3, and SIRT2, the DNA methyltransferase DNMT1, and the histone variant macroH2A^11–19^. Chromatin regulators that act as transcriptional activators, such as the histone acetyltransferases KAT1A and KAT2B, and the chromatin remodelers Brg1 and SRCAP have also been shown to regulate intestinal stem cell growth and differentiation^20–22^.

A potential role for factors involved in chromatin replication in the regulation of intestinal stem cells has not been explored in detail. Chromatin replication occurs during DNA replication to ensure proper packaging of the daughter duplexes. Parental nucleosomes are disrupted in front of the replisome and the resultant H3/H4 tetramers are recycled onto the newly replicated DNA behind the replication fork. The spatial retention of parental H3/H4 tetramers serves to maintain the context-dependent epigenetic memory of histone modification patterns^23–28^. The new histones are deposited via the replication-coupled chromatin assembly pathway. This pathway starts in the cytoplasm where newly synthesized H3/H4 dimers interact with the HAT1 complex that contains the histone acetyltransferase HAT1 and the histone chaperone Rbap46 (RBBP7). HAT1 acetylates the new H4 on K5 and K12, an evolutionarily conserved pattern that is specific for newly synthesized molecules^29–33^. The newly synthesized H3/H4 dimers are then transferred from the HAT1 complex to the histone chaperone ASF1, which facilitates nuclear import. Following import, the new H3 can also be acetylated, likely by CBP or KAT2A (GCN5)^34–45^. The new H3/H4 complexes are then directed to sites of replication and deposited by the chromatin assembly factor 1 (CAF1) complex^46–54^. The acetylation on the newly synthesized H3/H4 molecules is transient and is removed by the action of histone deacetylases, likely HDAC1 and HDAC2^55^. The deacetylation of the newly synthesized histones occurs over time with the removal of the new H4 di-acetylation (H4 K5/12ac) occurring over a roughly 2-hour period^29,56^. Maturation of the nascent chromatin is then completed by the transfer of the parental histone modification patterns to the new histones. Intriguingly, parental and newly-synthesized histones are asymmetrically distributed during the division of stem cells in the Drosophila intestine^57^.

We previously reported that murine Hat1 is essential for viability, as homozygous deletion of *Hat1* results in neonatal lethality caused by severe defects in lung development, leading to reduced aeration and respiratory distress, along with prominent craniofacial abnormalities of the skull and jaw^29^. At the cellular level, *Hat1*⁻^/^⁻ mouse embryonic fibroblasts display impaired proliferation, increased sensitivity to DNA-damaging agents, and marked genome instability^29^. Furthermore, haploinsufficiency of *Hat1* significantly decreases lifespan and produces an early-onset aging phenotype, including lordokyphosis, muscle atrophy, reduced subcutaneous fat, growth retardation, cancer, and paralysis, consistent with the observation that Hat1 expression declines in multiple tissues during normal aging in mice^58^.

The spectrum of phenotypes observed in HAT1 mouse mutants suggests that HAT1 may be involved in regulating the renewal or differentiation of stem cells. To explore this possibility, we have studied HAT1 in the context of intestinal stem cells (ISCs). We found that in the intestine, HAT1 is specifically expressed in the stem and progenitor cells in crypts. Inducible deletion of HAT1 in HAT1^fl/fl^; Villin-cre ERT2 mice results in a significant increase in crypt length with concomitant increases in proliferating cells. Differentiation is altered with skewing towards increased numbers of goblet cells and mis-localization of Paneth cells. Loss of HAT1 results in a dramatic loss of histone H4 K5 acetylation (H4K5ac) that is particularly pronounced in ISCs. Genome-wide mapping of H4K5ac indicates that HAT1-dpendent H4K5ac is localized to lamin-associated domains (LADs) and loss of H4K5ac is accompanied by increased levels of H3 K9 trimethylation (H3K9me3). Consistent with this observation, loss of HAT1 results in the decreased expression of multiple genes encoding for a-defensins, which are located in LADs. Importantly, HAT1 directly regulates ISC function, as it is required for the formation of intestinal organoids *ex vivo*.

## Materials and Methods

### Generation of *Hat1fl/fl* Mice with Villin-CreERT2 mediated intestinal deletion

Conditional *Hat1* floxed mice (*Hat1fl/fl*) were generated as previously described (*Nagarajan, et al.*, *PLOS Genetics*, 2013). Briefly, the targeting vector contained 5′ (6.9 kb) and 3′ (6.6 kb) homology arms, with two loxP sites flanking exon 3 of the *Hat1* gene and two FRT sites surrounding a *PGK*-*neo* cassette located downstream of exon 3. All fragments were generated by PCR using 129Sv/J genomic DNA and verified by restriction mapping and sequencing to confirm correct organization. The targeting vector was electroporated into Bruce4 embryonic stem (ES) cells, and correctly targeted clones were used to generate chimeric mice. Chimeras were crossed with C57BL/6J mice to obtain *Hat1+/fl* animals, which were intercrossed to produce the *Hat1fl/fl* line. To achieve tamoxifen-inducible, intestinal epithelium–specific deletion of *Hat1*, Hat1fl/fl mice were crossed with Villin-CreERT2 (Jax 020282) or Lgr5-EGFP-IRES-CreERT2 (Jax 008875) transgenic mice, generously provided by Maria Mihaylova. Cre-mediated recombination was induced by intraperitoneal (i.p.) injections of tamoxifen (Sigma-Aldrich, T5648) dissolved in corn oil (20 mg/mL) at 100 mg/kg body weight on alternating days for a total of five doses. Villin-CreERT2; Hat1fl/fl mice were used for broad intestinal epithelium–specific deletion, whereas Lgr5-EGFP-IRES-CreERT2; Hat1fl/fl mice allowed stem cell–specific deletion, as Lgr5 is expressed in stem cells at the base of the intestinal crypts. Both Cre lines were included in the control groups, which also comprised Hat1fl/fl mice lacking Cre and Cre-expressing mice carrying wild-type Hat1 alleles (Hat1+/+). Following the final tamoxifen injection, mice were maintained for 10 days to allow efficient recombination and epithelial turnover before euthanasia and tissue collection. Control mice received corn oil vehicles alone. All mice were maintained on a C57BL/6J background, and animals aged 3 and 18 months were used for experiments. All animal procedures were approved by the Institutional Animal Care and Use Committee (IACUC).

### Crypt Isolation and Culture of Intestinal organoids

Crypt isolation and organoid culture were performed as previously described^59^. Briefly, intestinal crypts were isolated from Hat1fl/fl, Hat1fl/fl-Villin-Cre and Hat1 fl/fl-LGR5-Cre mice following euthanasia. The duodenum and jejunum were excised, opened longitudinally, cleaned, and the intestinal epithelium was gently scraped using a glass slide. The collected tissue was washed repeatedly in PBS and incubated in PBS–EDTA for 40 min to release crypts. The crypt suspension was filtered through a 70 μm cell strainer and centrifuged at 250 × g for 5–10 min at 4 °C. The resulting crypt pellet was used for organoid culture. Crypts were resuspended in crypt culture medium consisting of Advanced DMEM/F 12 supplemented with Penicillin/Streptomycin, L-Glutamine, 200 ng/ml Noggin (Peprotech), 50ng/ml R-spondin, 1X B27 (Life Technologies), 3 μM CHIR99021 (Stem Cell Technologies), 1X N2 (Life Technologies), 1 μM Nacetyl-L-cysteine (Sigma Aldrich),10 μM Y-27632 dihydrochloride monohydrate (Sigma Aldrich), and 15 ng/ml EGF (Peprotech). The suspension was examined under a microscope to confirm crypt density and mixed with Matrigel at a 30:70 ratio. Droplets (25 μL) were plated in 48-well plates, incubated at 37°C for 15 minutes to polymerize, and 250 μL of medium was added per well. Medium was refreshed every 2–3 days. For in vitro deletion of *Hat1* in intestinal organoids, Tamoxifen was added to the culture medium at a final concentration of 0.5–1 µM. For passaging, Matrigel domes were disrupted by pipetting and scraping, collected, and centrifuged at 250–300 × g for 5 min at 4°C. Organoids were resuspended in fresh medium, mixed with Matrigel (30:70), plated as 25 μL droplets, and incubated at 37°C for 15 min before adding additional medium.

### Whole-Mount Organoid Immunostaining

Organoids were washed with PBS and fixed in freshly prepared 4% paraformaldehyde at room temperature for 20 min. Following fixation, organoids were collected in 1.5 mL tubes using PBS containing 1% BSA to prevent adhesion, centrifuged at 250 × g for 5 min, and washed once with PBS. Permeabilization and blocking were performed in 0.2% Triton X-100 and 2% normal goat serum in PBS (PBST) for 30 min at room temperature, followed by PBST washes. Organoids were incubated with primary antibodies (Hat1, LGR5, GFP) diluted in PBST with 2% normal goat serum overnight at 4°C, washed with PBST, and then incubated with secondary antibodies (anti rabbit; anti chicken) under the same conditions for 1 h at room temperature. Nuclei were counterstained with DAPI for 5 min, washed, resuspended in mounting medium, and distributed onto glass slides. Coverslips were applied, and slides were allowed to cure for ∼30 min before imaging. Imaging was taken under a confocal microscope.

### Western Blot Analysis

Whole-protein lysates from intestinal crypt cells were prepared using RIPA buffer (Research Products International, R26200; 100 mM Tris-HCl, pH 7.4; 300 mM NaCl; 2% NP-40; 1% sodium deoxycholate; 0.2% SDS). Equal amounts of protein were resolved on 10% or 18% SDS–polyacrylamide gels and transferred to nitrocellulose membranes (GE Healthcare Life Sciences, Cat. No. 10600004). Membranes were blocked for 1 hour at room temperature in 5% skim milk prepared in TBS-T (20 mM Tris-HCl, pH 7.4; 150 mM NaCl; 0.1% Tween-20) and then incubated overnight at 4°C with primary antibodies against Hat1 (Abcam, ab12163), β-Actin (Abcam, ab8227), H3 (Abcam, ab1791), H4K5 (Abcam, ab51997), and H4K12 (Abcam, ab46983). Following incubation with HRP-conjugated secondary antibodies, protein bands were visualized using the iBright Imaging System (Invitrogen).

### Histology and Immunohistochemistry

Mouse intestinal tissues were fixed in 10% neutral buffered formalin phosphate solution, embedded in paraffin, and sectioned at a thickness of 5-8µm. For histological assessment, sections were stained with hematoxylin and eosin (H&E) for general morphology and Alcian blue for goblet cell visualization. For immunohistochemistry, antigen retrieval was performed in Target Retrieval Solution (Dako North America, Inc., Cat. No. S1700) using a pressure cooker (Instant Pot) for 20 minutes, followed by incubation in 0.3% hydrogen peroxide for 30 minutes at room temperature to quench endogenous peroxidase activity. Slides were then blocked with 5% normal donkey serum in TBST for 30 minutes, followed by 15-minute incubations in 5% donkey serum containing avidin/biotin blocking solutions (Vector Laboratories). Primary antibodies included OLFM4 (CST-39141) and Cleaved Caspase-3 (CST-9664). A biotin-conjugated donkey anti-rabbit secondary antibody (Jackson ImmunoResearch) was applied, followed by detection with the VECTASTAIN® Elite ABC Kit (Vector Laboratories) and visualization using Dako Liquid DAB+ Substrate (Dako). Sections were counterstained with hematoxylin and mounted with Cytoseal.

### Confocal microscope Immunofluorescence (IF) analysis

Mouse intestinal tissues were fixed in 10% buffered formalin phosphate solution (Fisher Scientific, Cat. No. SF100-4) for 48 hours and subsequently transferred to 70% ethanol. Fixed tissues were processed, embedded in paraffin, and sectioned at a thickness of 5 µm onto positively charged glass slides. Paraffin sections were deparaffinized through a xylene–ethanol series and subjected to antigen retrieval using Target Retrieval Solution (Dako North America, Inc., Cat. No. S1700) in a steamer for 60 minutes. After cooling to room temperature (RT), sections were blocked for 60 minutes in 1% antibody dilution buffer (10% ADB in PBS, pH 7.4; containing 3% BSA, 10% goat serum, and 0.05% Triton X-100). Slides were then incubated overnight at 4°C with primary antibodies against Hat1 (Abcam, ab12163), H4K5 (Abcam, ab51997), LGR5 (Thermo-MA5-25644) anti-GFP (Aves Labs, GFP-1020), Ki67 (CST #9129), DCLK1 (CST #62257), Lysozyme-LYZ (Thermo RB-372-A1) and ChgA (Santa Cruz, sc-1488), diluted in antibody dilution buffer. The following day, sections were washed three times with 1% ADB and incubated for 60 minutes at RT in the dark with Alexa Fluor® 594 and 488–conjugated secondary antibody (Jackson ImmunoResearch Laboratories, Inc.) diluted in ADB. After three washes with PBS, slides were mounted with VECTASHIELD® mounting medium containing DAPI (Vector Laboratories, Cat. No. H1200). Fluorescent images were acquired using a confocal microscope and analyzed with ImageJ software.

### RNA Extraction and Digital droplet PCR (ddPCR)

Total RNA was extracted from mouse intestinal crypt cells using the NucleoSpin RNA Plus Kit (Takara Bio). cDNA was synthesized from 500 ng of RNA using the High-Capacity cDNA Reverse Transcription Kit (Applied Biosystems). *Hat1* mRNA expression was quantified using droplet digital PCR (Bio-Rad) following the manufacturer’s protocol. Data were analyzed with QuantaSoft software using Poisson statistics, and expression levels were normalized to *Cyclophilin* mRNA. Primer and probe sequences were: *Hat1* (Forward 5′-CTGAGCAATACAGAAGCTACAG-3′; Reverse 5′-TCTGGTCTCAGGCATTTCTTC-3′; Probe FAM-5′-ACAAGAAAAAGCAGAGGGATCTTGCCAAGA-ZEN/IBFQ-3′) and *Cyclophilin* (Forward 5′-GTCAACCCCACCGTGTTCTT-3′; Reverse 5′-TTGGAACTTTGTCTGCAAACA-3′; Probe VIC-5′-CTTGGGCCGCGTCT-MGB-3′).

### CUT&Tag

CUT&Tag for H3K9me3 (Abcam, ab8898) and H4K5ac (Abcam, ab51997) was performed on isolated intestinal crypt cells as described in Martin et al. (2025) based on the Epicypher CUT&Tag protocol. Higher than recommended cell numbers (>250,000 cells) were used for each reaction to improve yields. Experiments were performed in biological duplicate, with cells isolated from two different mice for each condition (WT and Hat1-deleted), except for H4K5ac where 3 mice were obtained.

### CUT&Tag Data Processing and Analysis

CUT&Tag data was processed as described in Martin et al. (2025) to generate bigwig files which were visualized in Integrative Genomics Viewer (IGV). The H4K5ac CUT&Tag reads was trimmed before aligning using TrimGalore! (version 0.6.10) to improve alignment by bowtie2^60^. The H3K9me3 CUT&Tag reads aligned efficiently without trimming. Peak calling was done using SEACR using a threshold of 0.01 under the “stringent” setting without normalization. Differential peak analysis was performed with the dIffbind package in R and plotted using ggplot2^61^.

### RNA Sequencing

Total RNA was extracted from intestinal crypts using the NucleoSpin RNA Plus Kit (Takara Bio) according to the manufacturer’s instructions. RNA quality was assessed with an Agilent 2100 Bioanalyzer, and libraries were prepared from 200 ng of total RNA using the TruSeq RNA Sample Prep Kit v2 (Illumina). The concentration and size distribution of the completed libraries were verified using an Agilent Bioanalyzer. Libraries were sequenced to a depth of 37–45 million pass-filter clusters per sample following standard Illumina protocols with the cBot and cBot Paired-End Cluster Kit v3. Sequencing was performed on an Illumina HiSeq 2000 using the TruSeq Sequencing Kit and HCS v2.0.12 data collection software. Transcriptome analysis of HAT1^Villin WT^ and HAT1^Villin KO^ organoids was carried out in biological triplicates, and statistical analyses were conducted in R. However, one sample of HAT1^Villijn KO^ was eliminated from the analysis due to degradation.

### RNA-seq Data Processing and Analysis

RNA sequencing data was processed using the nf-core RNA-seq pipeline (version 3.13.2) with quantification by Salmon and the output was analyzed in R. Differential gene expression analysis was carried out with DESeq2 v1.34.0 and a q-value threshold of 0.05 was used to determine which genes were significantly differentially expressed^62^. For the heatmap, each row was independently converted to z-scores and plotted using the pheatmap package, v1.0.13 (https://cran.rproject.org/web/packages/pheatmap/index.html) with default row clustering and no column clustering. Gene Ontology Enrichment analysis was performed with the enrichr package, v3.2 (https://cran.r-project.org/web/packages/enrichR/index.html). Other plots were generated using ggplot2.

### Statistical Analysis

Statistical analyses were performed using either SigmaPlot version 12.0 (Systat Software) or in R using using an unpaired Student’s t-test ir a Wicoxen’s rank sum test. Data are presented as mean ± standard error (SE). Statistical comparisons between experimental groups were conducted using an unpaired Student’s *t*-test, and corresponding *p*-values are indicated in the graphs. Unless otherwise stated, all experiments were independently repeated at least three times. *N* represents biological replicates. Age- and sex-matched mice (3 and 18 months old) were randomly assigned to experimental groups. All bar plots were generated in R using ggplot2 except for those in Figures 3, 7, and S2, which were made in Excel.

## Results

### HAT1 is required for the maintenance of intestinal crypt structure

Epithelial cells of the intestine are organized along the crypt-villus axis. At the base of the villus is a structure known as the crypt that is the location of the stem cells and their niche. As the stem cells divide, cells that will differentiate move up the crypt into the transit amplifying zone where they become progenitor cells. The intestinal stem and progenitor cells differentiate into 2 lineages: the secretory and absorptive. The secretory lineage includes Paneth, goblet, tuft and enteroendocrine cells, while the absorptive lineage is comprised primarily of enterocytes. Cells become fully differentiated as they leave the transit amplifying zone and move into the villus^1^. Immunofluorescent staining of mouse proximal small intestine sections with α-HAT1 antibodies indicates that HAT1 is selectively expressed in intestinal crypts, in both the base of the crypt and in the transit amplifying zone Figure 1A). To confirm that HAT1 is expressed in ISCs, we stained sections of the proximal small intestine from mice expressing GFP from the endogenous Lgr5 promoter with α-HAT1 antibodies^63^. HAT1 is expressed in both the Lgr5-positive cells at the base of the crypt and progenitor cells in transit amplifying zone (Figure 1B). Interestingly, there little or no expression of HAT1 in the differentiated cells in the villi or in the Paneth cells in the crypt base.

**Figure 1.**
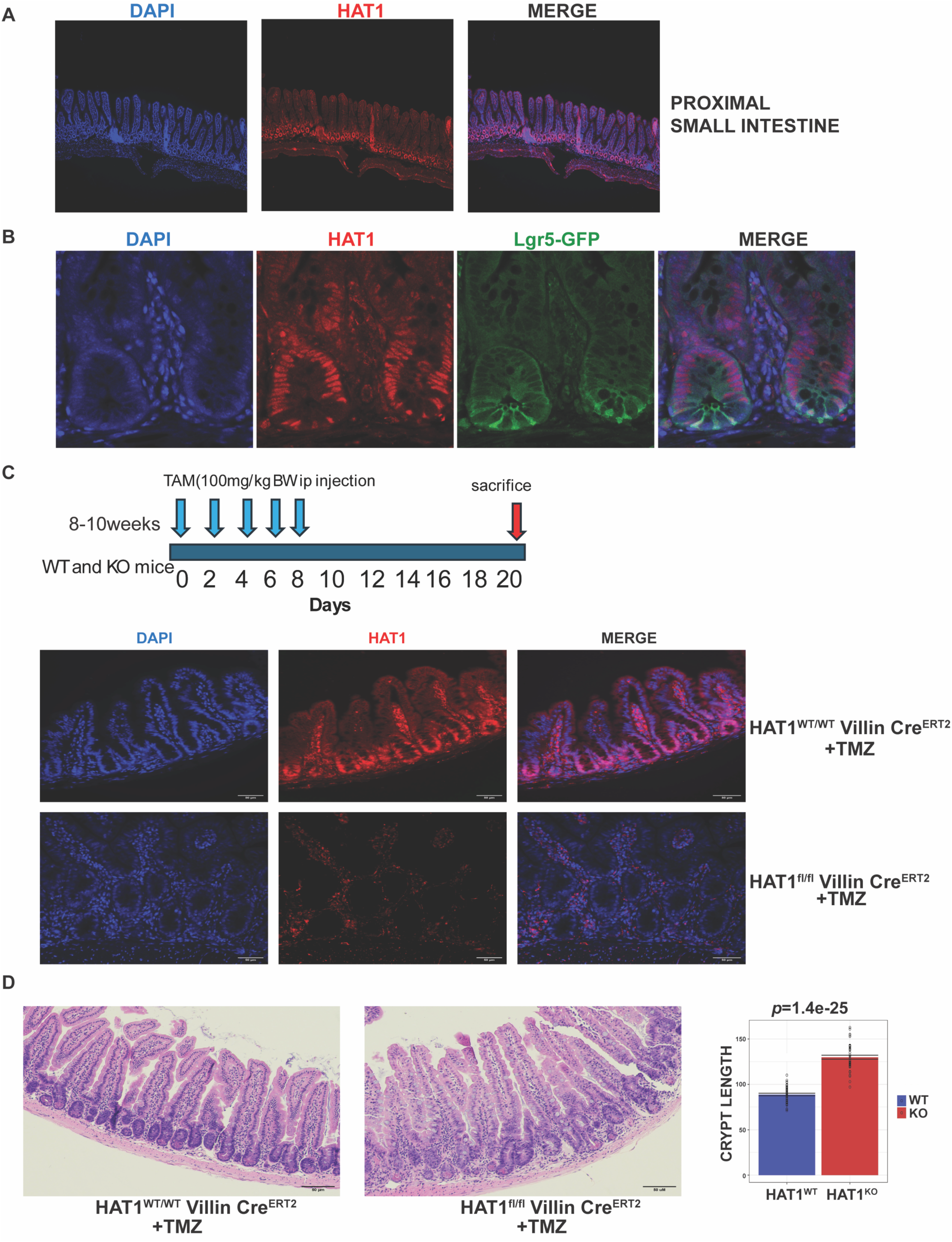
HAT1 Is Essential for Intestinal Crypt Maintenance. **(A)** Representative immunofluorescent confocal images of proximal small intestine sections from WT mice stained with α-HAT1 antibodies. (N = 3-5 mice per group) **(B)** Immunofluorescent confocal images of proximal small intestine sections from Lgr5–GFP mice stained with α-HAT1 antibodies, allowing comparison of HAT1 expression with Lgr5-positive intestinal stem cells. **(C)** Schematic diagram of the tamoxifen-inducible Villin-Cre model. Three-month-old HAT1^villin WT^ and HAT1^Villin KO^ mice were administered Tamoxifen once daily for 5 days to induce HAT1 deletion and were sacrificed 10 days after the final injection. Immunofluorescent staining of small intestine sections confirms efficient HAT1 loss in HAT1^Villin KO^ mice compared with WT controls. **(D)** Representative H&E-stained sections of proximal small intestine from HAT1^villin WT^ and HAT1^VILLIN KO^ mice, with quantification of crypt length (n=250 crypts, N = 3 mice per group). All images include a 50 μm scale bar.

To determine whether HAT1 is functionally important for intestinal function, we generated a HAT1^fl/fl^; Villin-Cre^ERT2^ mouse model, which allows for the tamoxifen-inducible deletion of HAT1 in the epithelial cells of the intestine^64^. These mice will be referred to as HAT1^villin WT^ and HAT1^villin KO^. At 3 months of age, HAT1^villin WT^ and HAT1^Villin KO^ mice were injected with tamoxifen once per day for 5 days and sacrificed after 10 days (Figure 1C). PCR, Western blot, and immunofluorescence analyses show that HAT1 is efficiently deleted (Figure 1C, Supplementary Figure 1A and 1B). Loss of HAT1 had a minor effect on body weight and no significant effect on overall small intestine or colon length (Supplementary Figure 1C). Morphological analysis of sections from the proximal small intestine reveals a dramatic effect of HAT1 loss on the structure of the intestinal crypt (Figure 1D). The length of the crypt becomes significantly elongated and the regular architecture of the ISC and Paneth cells at the base of the crypt is disrupted.

### HAT1 restricts the proliferation of intestinal stem and progenitor cells

To determine whether the alterations in the intestinal crypts are the result of increased intestinal stem and progenitor cell proliferation, we stained tissue sections for Ki67. Figure 2A shows a striking increase in cellular proliferation in the crypt, with elongated crypts populated by Ki67+ cells. We then stained sections for the ISC marker OLFM4 to test whether the increase in proliferative cells in the crypts of HAT1^villin KO^ mice following tamoxifen treatment is accompanied by an increase in the number of ISCs and early progenitor cells. As seen in Figure 2B, there is an ∼2-fold increase in the number of OLFM4 positive cells following the loss of HAT1. One explanation for this observation is that there is a decrease in apoptosis after HAT1 loss. This was tested by staining the sections for cleaved caspase 3. The results indicate that there is an increase in apoptosis in the absence of HAT1 (Figure 2C). These results suggest that HAT1 normally functions to restrict the proliferation of intestinal stem and progenitor cells.

**Figure 2.**
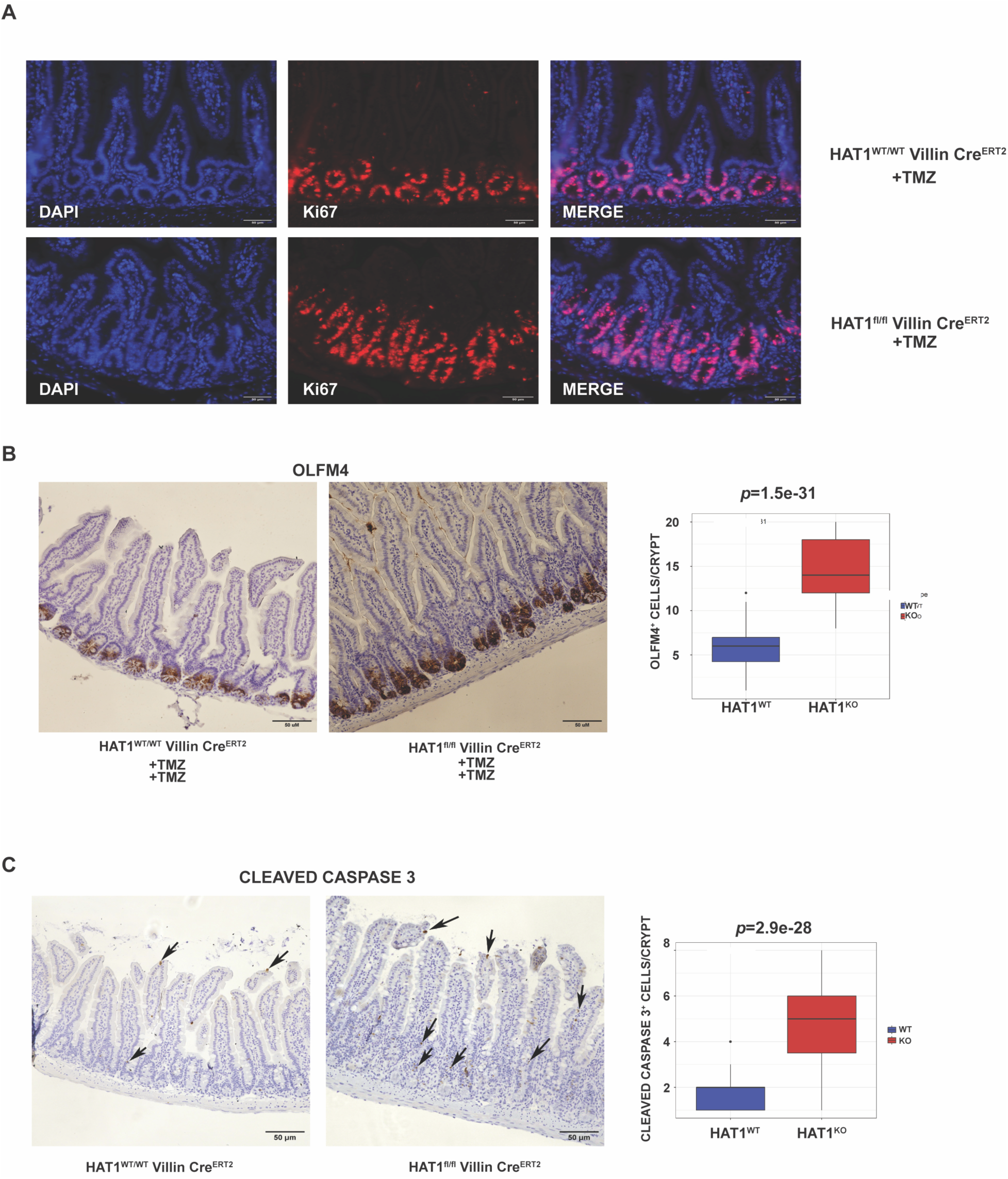
HAT1 Restrains Intestinal Stem and Progenitor Cell Expansion. **(A)** Representative confocal images of proximal small intestine sections from HAT1^villin WT^ and HAT1^Villin KO^ mice stained with Ki67, a marker of proliferating cells in the crypts. **(B)** Representative immunohistochemistry (IHC) images of proximal small intestine sections from HAT1^villin WT^ and HAT1^Villin KO^ mice stained for OLFM4, an intestinal stem cell marker. Quantification of OLFM4-positive cells per crypt is shown. **(C)** Representative immunohistochemistry IHC images of proximal small intestine sections HAT1^villin WT^ and HAT1^Villin KO^ mice stained for cleaved caspase-3, an apoptotic marker. Quantification of cleaved caspase-3–positive cells per crypt is shown. (N = 3 mice per group). All images include a 50 μm scale bar

Alternatively, if HAT1 is required for ISC viability, loss of HAT1 may be triggering a regenerative response, leading to increased ISC proliferation.

### HAT1 regulates ISC differentiation

While HAT1 clearly regulates ISC proliferation, we next asked whether HAT1 is also involved in in differentiation. We first stained proximal small intestine sections with Alcian Blue to visualize and quantify goblet cells. There is a significant increase in the number of goblet cells following deletion of HAT1 and many of these goblet cells display an altered morphology (Figure 3A). Paneth cells were visualized by staining for LYZ1 (Figure 3B). When HAT1 is deleted from 3-month-old animals using the strategy depicted in Figure 1C (short-term deletion), there is little effect of HAT1 loss on Paneth cells (Figure 3B). However, if the HAT1 deletion is maintained for 18 months through weekly tamoxifen administration (long-term deletion), there is a dramatic redistribution of Paneth cells in the HAT1^Villin KO^ intestines, with LYZ1 positive cells found throughout the villi. DLCK1 staining indicates that there is small increase in tuft cells following HAT1 loss in the intestine (Figure 3C) and a similar increase in enteroendocrine cells, as indicated by CHGA staining (Figure 3D). The alterations in cells of the secretory lineage indicate that HAT1 is required to maintain normal ISC differentiation *in vivo*.

**Figure 3.**
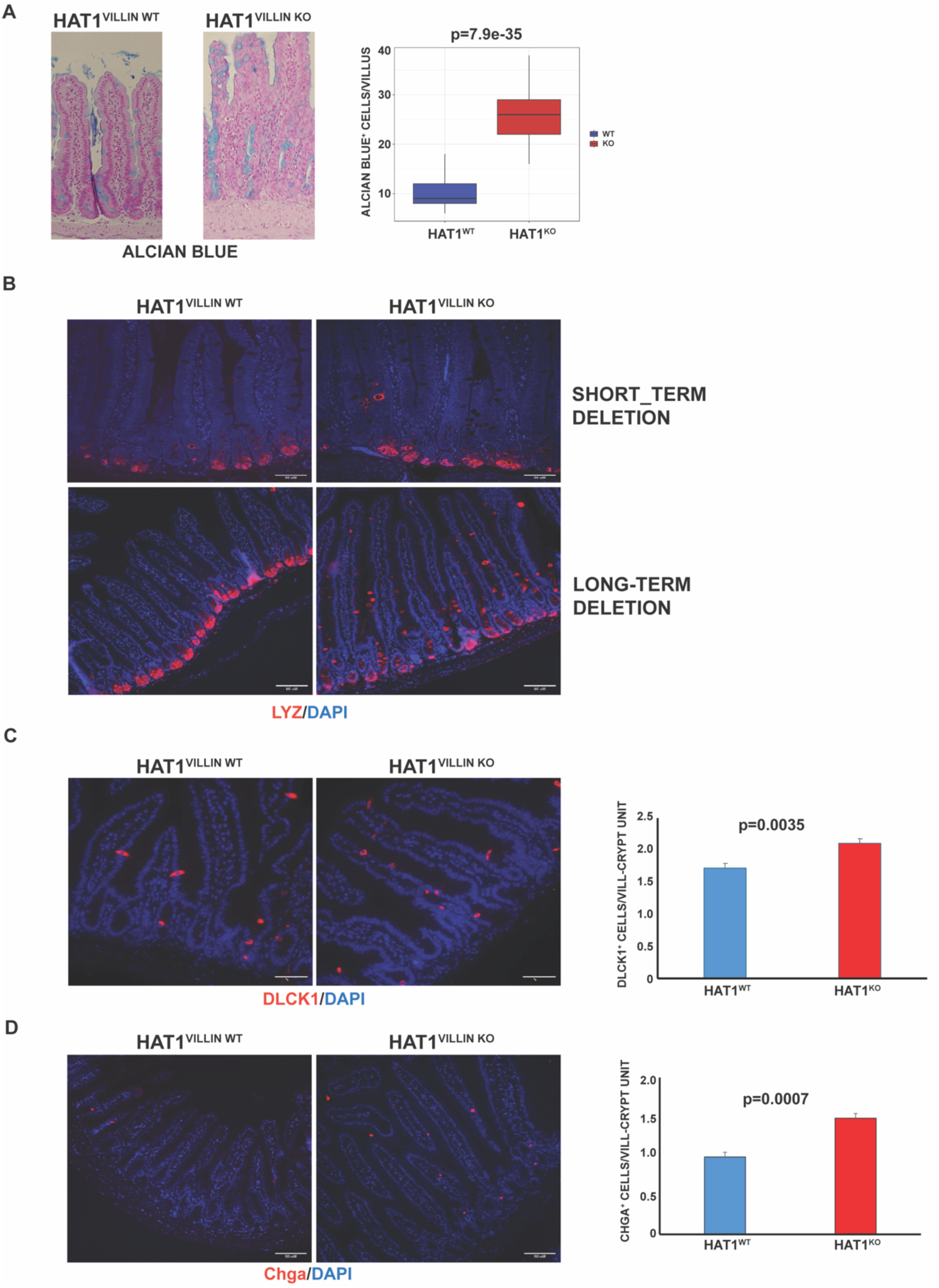
HAT1 Regulates Intestinal Stem Cell Differentiation. **(A)** Representative Alcian Blue–stained sections of proximal small intestine from HAT1^villin WT^ and HAT1^Villin KO^ mice, highlighting goblet cells. Quantification of Alcian Blue–positive cells per villus is shown (N = 3 mice per group). **(B)** Representative confocal images of proximal small intestine sections from HAT1^villin WT^ and HAT1^Villin KO^ mice stained with lysozyme (LYZ), a Paneth cell marker, comparing Paneth cell distribution following short-term and long-term tamoxifen-induced HAT1 deletion. **(C)** Confocal immunofluorescence of proximal small intestine sections from HAT1^villin WT^ and HAT1^Villin KO^ mice stained with DCLK1, a tuft cell marker, showing tuft cell localization and abundance in crypts and villi. Quantification of DCLK1-positive cells per crypt and per villus is shown (N = 3 mice per group). **(D)** Representative confocal images of proximal small intestine sections from HAT1^Villin WT^ and HAT1^Villin KO^ mice stained with chromogranin A (ChgA), a marker of enteroendocrine cells. Quantification of ChgA-positive cells per crypt and per villus is shown (N = 3 mice per group). All images include a 50 μm scale bar

### HAT1 is required for H4K5ac in ISCs

HAT1 was originally isolated based on its ability to acetylate newly synthesized (and non-nucleosomal) histone H4 on lysine residues 5 and 12^30^. To determine whether HAT1 is responsible for H4K5ac and H4K12ac in the intestine, we analyzed extracts derived from crypts from proximal small intestine isolated from HAT1^villin WT^ and HAT1^Villin KO^ animals. We observed a significant decrease in H4K5ac in the HAT1^VILLIN KO^ crypt extracts but no change in the levels of H4K12ac (Figure 4A). We performed further analysis using immunofluorescence to localize HAT1-dependent H4 acetylation in the intestine. As seen in Figure 4B, H4K5ac is largely limited to cells of the intestinal crypt, with H4K5ac more prominent in ISCs relative to progenitor cells in the transit amplifying zone. Following loss of HAT1, there is a decrease in H4K5ac levels in crypts, which is particularly pronounced in ISCs (Figure 4B, lower panel).

**Figure 4.**
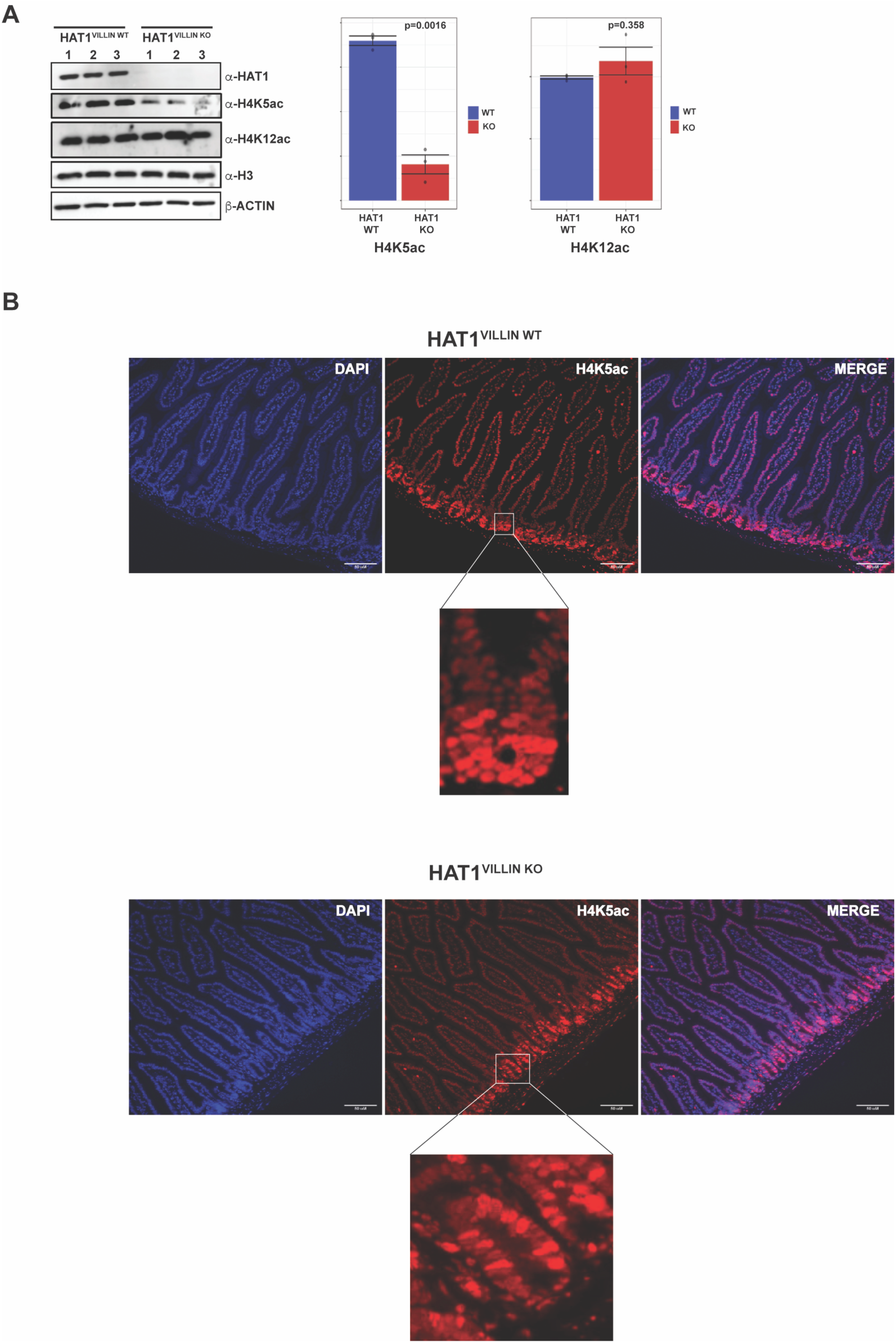
HAT1 Is Essential for Histone H4K5 Acetylation in ISCs. **(A)** Representative Western blot analysis of proximal small intestine crypt protein lysates from three HAT1^villin WT^ and three HAT1^Villin KO^ mice. Blots were probed for HAT1, H4K5ac and H4K12ac, total H3, and Actin as a loading control. The accompanying histogram shows quantification of H4K5ac and H4K12ac levels normalized to total H3 (N = 3 mice per group). **(B)** Representative confocal images of proximal small intestine sections from HAT1^Villin WT^ and HAT1^Villin KO^ mice stained for H4K5ac to visualize histone acetylation in the crypts. The crypt region is shown at higher magnification in an inset. Images are representative of N = 3 mice per group. All images include a 50 μm scale bar.

H4K12ac levels are also enriched in crypts in the proximal small intestine, similar to the pattern for H4K5ac. However, consistent with the Western blot data, there is no change in intestinal H4K12ac staining following the loss of HAT1 (Figure S3).

### HAT1 regulates histone modification patterns in lamin-associated domains

We determined the genome-wide localization of H4K5ac in cells isolated from proximal small intestine crypts using CUT&Tag^65^. Following peak calling, we looked for differences in H4K5ac peaks in crypts isolated from HAT1^VILLIN WT^ and HAT1^VILLIN KO^ animals. Surprisingly, there were no significant changes in H4K5ac peaks following loss of HAT1 (Figure 5A). We plotted the log_2_ Fold change of H4K5ac in HAT1^VILLIN KO^ relative to HAT1^VILLIN WT^ across the genome. As seen in Figure 5B, rather than decreases in individual peaks, H4K5ac decreases in HAT1^VILLIN KO^ across large domains. This broad loss of H4K5ac is easily observed in a browser track by overlaying the H4K5ac signal from HAT1^VILLIN KO^ and HAT1^VILLIN WT^ crypt cells (Figure 5B). The domains that show a HAT1-dependent loss of H4K5ac contain relatively low levels of this acetylation suggesting that they are heterochromatic. Indeed, these domains display a high degree of overlap with the locations of lamin-associated domains (LADs) in NIH3T3 cells previously observed (Figure 5B)^66^.

**Figure 5.**
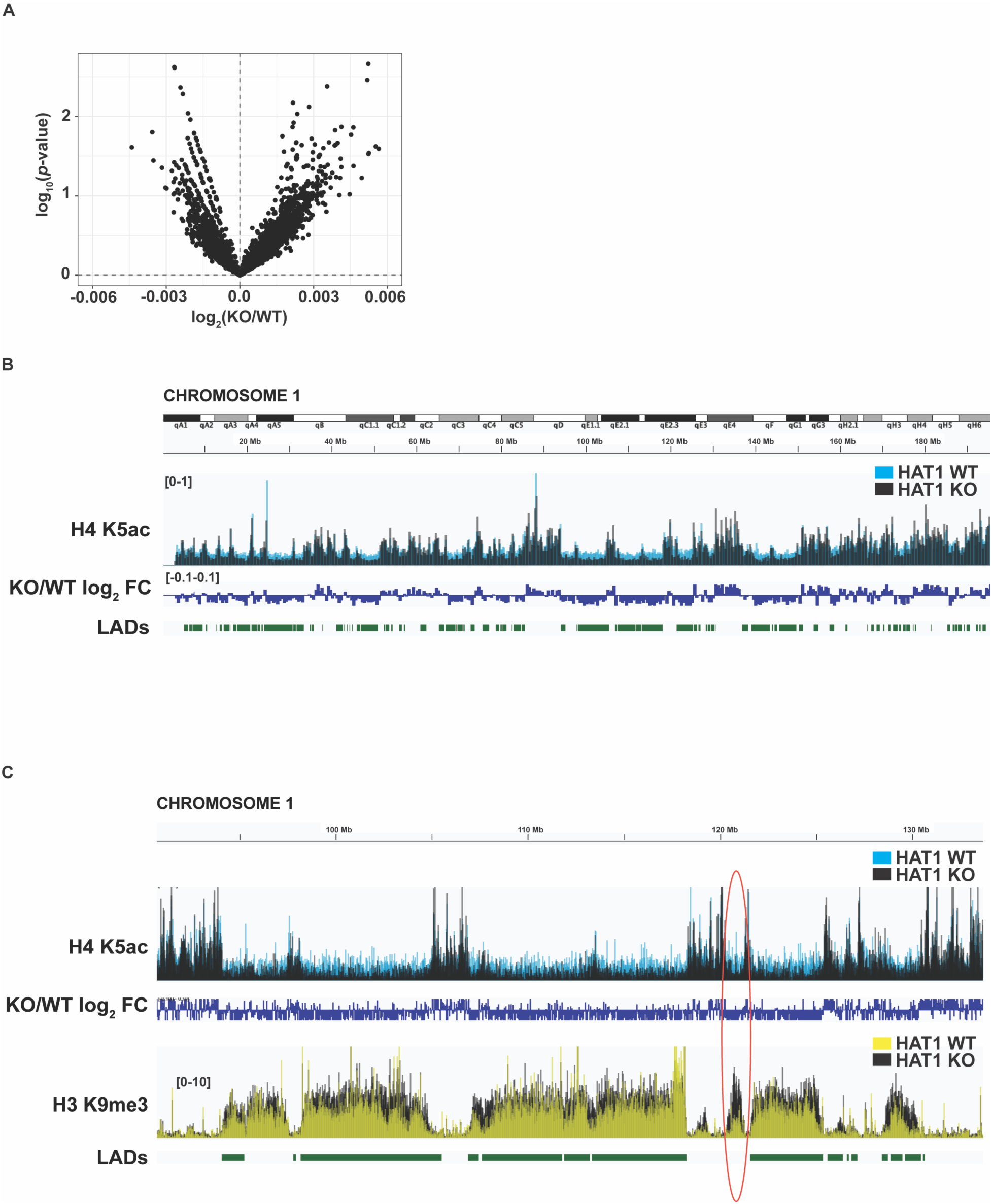
HAT1 Regulates the Structure of Lamin-Associated Domain Chromatin. **(A)** Volcano plot of H4K5ac peaks in HAT1 WT vs KO crypts. **(B)** IGV Browser view illustrating changes in H4K5ac in intestinal crypt cells upon HAT1 deletion. **(C)** IGV Browser view comparing changes in H4K5ac and changes in H3K9me3 on chromosome 1.

A specific effect of HAT1 in LADs is consistent with a recent study that showed that HAT1 is essential for maintaining chromatin accessibility in domains termed HAT1-dependnet accessibility domains (HADs) that overlap with a subset of LADs in mouse embryonic fibroblasts (MEFs). Loss of HAT1 also resulted in an increase in the density of H3K9me3 in these HADs/LADs^67^. To determine whether HAT1 also regulates H3K9me3 in intestinal crypts, we used CUT&Tag to map this modification. Analysis of the H3K9me3 signal shows that intestinal crypts contain broad domains of H3K9me3 that colocalize with most of the LADs that have previously been mapped in MEFs (Figure 5C). In addition, intestinal crypts have domains of H3K9me3 that do not coincide with LADs in MEFs (an example is denoted by the red oval in Figure 5C). This indicates that the LAD structure in intestinal crypt cells is likely to be similar that observed in MEFs but that there is a subset of LADs that are unique to thes cell types.

Importantly, LADs that show a HAT1-dependent decrease in H4K5ac show a concomitant increase in H3K9me3. This suggests that rather than regulating specific peaks of H4K5ac, HAT1 regulates multiple aspects of the chromatin structure of LADs in intestinal crypt cells.

### HAT1 regulates the expression of α-defensin genes

Gene expression in intestinal crypt cells from HAT1^VILLIN WT^ and HAT1^VILLIN KO^ animals was analyzed by RNA-Seq. Short term loss of HAT1 had a small effect on transcript levels with 144 genes down-regulated and 235 genes up-regulated (Figure 6A, Table S1). KEGG pathway analysis of the differential expressed genes identifies several pathways that are mis-regulated. The most significant was *Staphylococcus aureus* infection. Examination of the genes that make up this pathway with the differentially expressed genes in HAT1^VILLIN KO^ crypts highlights the family of α-defensin genes^68^. There are 19 α-defensin genes that are down-regulated in HAT1^VILLIN KO^ crypts (Figure 6C). The α-defensin gene are also among the most highly abundant of the differentially expressed genes in the intestinal crypts (Figure 6D, Table S1). The high level of α-defensin gene expression is likely from mature Paneth cells or secretory precursor cells^68^. The α-defensins are in a large cluster on chromosome 8. Interestingly, this cluster is located in two adjacent LADs, which display a significant decrease in H4K5ac in HAT1^VILLIN KO^ crypts (Figure 6E). Together, these results indicate that HAT1 can influence gene expression in cells of the intestinal crypt and that the regulation of LAD chromatin structure by HAT1 may be involved in this process. However, HAT1 may also influence ISC proliferation and differentiation by additional mechanisms.

**Figure 6.**
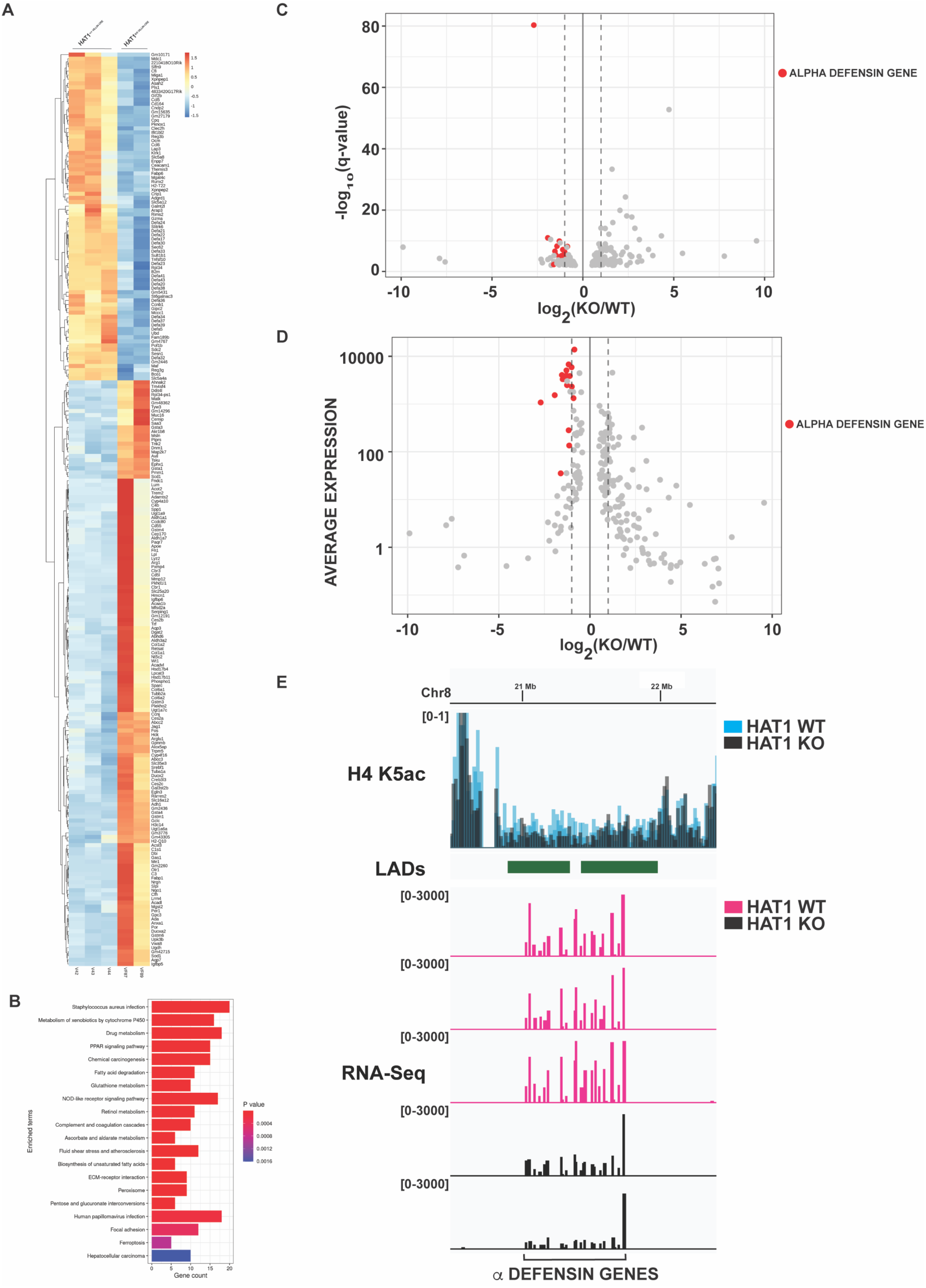
HAT1 Regulates the Expression of α-defensins. **(A)** Hierarchical clustering heatmap of differentially expressed genes (q-value<0.01). **(B)** Terms from the “KEGG 2019 Mouse” database that are enriched in differential genes (q-value<0.05). **(C)** Volcano plot of differentially expressed genes (q-value<0.01). **(D)** Average expression across all WT and KO replicates plotted against log2-fold change for differentially expressed genes (q<0.01). **(E)** Illustration of LAD with downregulation of α-defensin genes alongside loss of H4K5ac.

### HAT1 is required for ISC function and differentiation

To determine whether the defects in intestinal crypts observed following HAT1 loss are a direct result altered ISC function, we exploited the intestinal organoid model system^69^. Crypts were isolated from HAT1^Villin WT^ and HAT1^Villin KO^ mice following tamoxifen injection and plated in Matrigel with specific crypt media necessary for the development of intestinal organoids^59,70^. Normal organoids formed from cells isolated from the HAT1^Villin WT^ mice, while very few organoids were generated from crypt domains derived from HAT1^Villin KO^ mice (Figure 7A). To test the necessity of HAT1 in organoid regeneration and confirm that ISCs from the HAT1^Villin KO^ mice were competent to form intestinal organoids, we isolated crypts from non-tamoxifen induced animals and generated intestinal organoids first. The organoids were then dispersed into fresh media and 4-OH tamoxifen was added to one set of wells to delete the *HAT1* gene *ex vivo*. Strikingly, deletion of HAT1 in established organoids completely disrupted the ability of the ISCs to differentiate and we primarily observed spherical structures reminiscent of enterocysts (Figure 7B)^71^.

**Figure 7:**
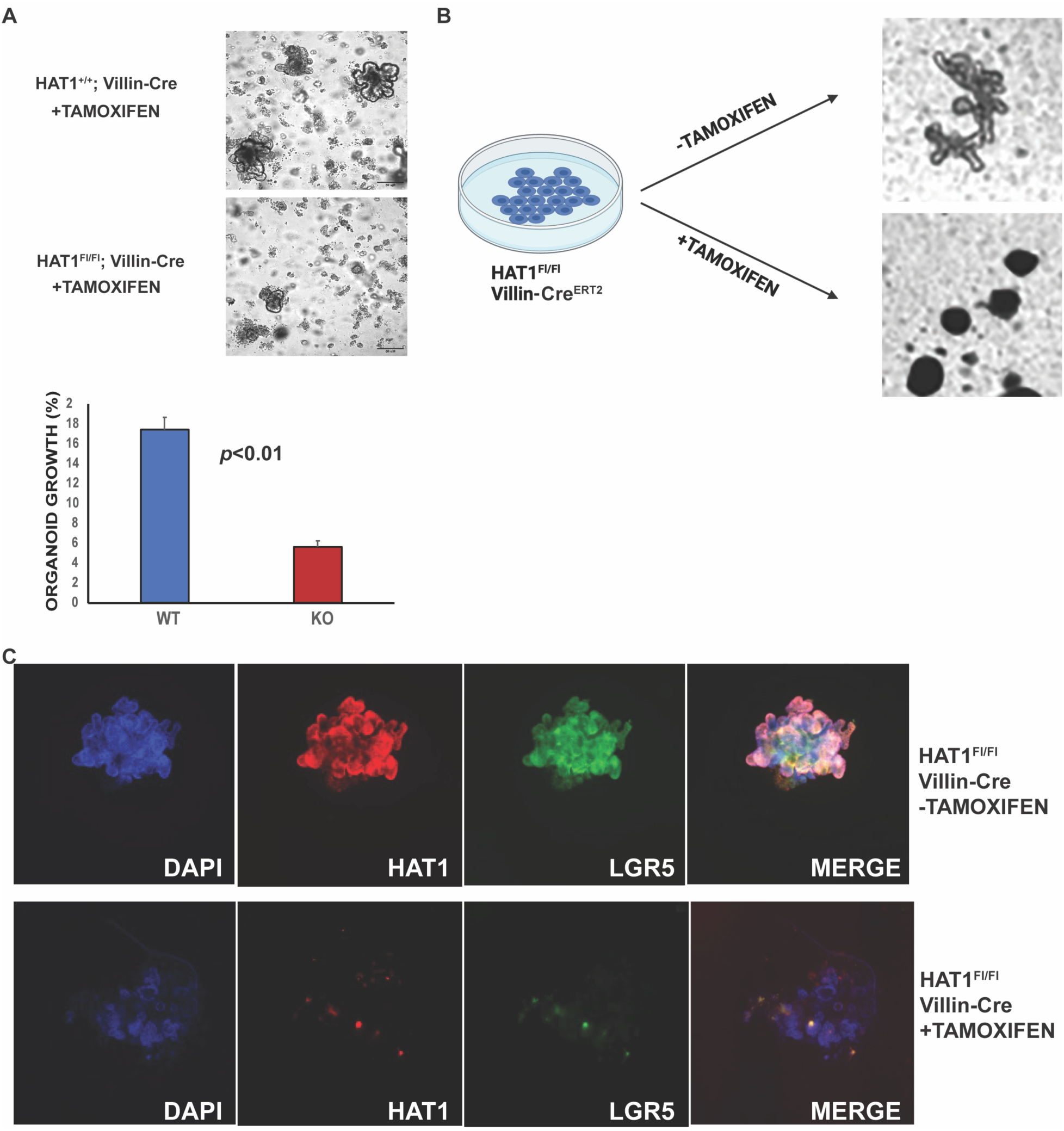
HAT1 Is Essential for Intestinal Stem Cell Function and Differentiation. **(A)** Representative images of intestinal organoids derived from crypts isolated from HAT1^villin WT^ and three HAT1^Villin KO^ mice following tamoxifen injection. Crypts were plated in Matrigel with specialized crypt media to support organoid growth. The accompanying histogram shows the percentage of organoid formation comparing WT and KO (N = 5 mice per group). P < 0.01. **(B)** Crypts from HAT1^fl/fl^ Villin-Cre mice without prior *in vivo* tamoxifen treatment were cultured to generate intestinal organoids. 4-Hydroxytamoxifen (4-OHT) was added to one set of wells to induce HAT1 deletion *in vitro*, while another set remained untreated (–4-OHT). Representative images show the effect of HAT1 deletion on organoid development. Scale bars, 50 μm. **(C)** Whole-mount immunofluorescence of intestinal organoids derived from HAT1^fl/fl^ Villin Cre crypts cultured without (–TAM) or with (+TAM) 4-hydroxytamoxifen to induce HAT1 deletion *in vitro*. Organoids were stained for HAT1 and LGR5 to assess HAT1 expression and intestinal stem cell marker localization. Scale bars, 50 μm.

To directly test whether HAT1 is required for the maintenance of ISC in organoid cultures, we stained organoids for HAT1 and LGR5 (Figure 7C). In the absence of tamoxifen treatment, organoids from HAT1^fl/fl^ VILLIN-Cre mice indicate that HAT1 is enriched in cells that express high levels of Lgr5. However, following tamoxifen treatment, there is an almost complete lack of both HAT1 and Lgr5 expression. Strikingly, there several cells that have escaped HAT1 deletion and these cells retaining Lgr5 expression. Combined, these studies demonstrate that HAT1 plays a key role in the regulation of ISC self-renewal, proliferation and differentiation.

## DISCUSSION

Our results indicate that HAT1 plays a critical role in regulating the proliferation and differentiation of ISCs. Intestine-specific loss of HAT1 results in alterations in intestinal architecture with elongated crypts resulting from increased proliferation of ISCs. HAT1 loss also results in significant alterations in secretory cells, and more specifically Goblet and Paneth cell populations, where the number of Goblet cells increases and cells expressing Paneth cell markers mislocalize in villi. At the molecular level, HAT1 loss has a significant effect on the structure of chromatin in cells of the intestinal crypt. There is a dramatic loss of H4K5ac, particularly in ISCs, that is reflected in a decrease in this acetylation throughout LADs. This loss of H4K5ac in LADs is accompanied by an increase in H3K9me3. A direct role for HAT1 in ISC function is clearly demonstrated by the requirement for HAT1 for the generation of intestinal organoids *in vitro*.

Several histone modifying enzymes have been shown to be involved in regulating the maintenance or differentiation of ISCs. This includes the histone acetyltransferases KAT2A/KAT2B (GCN5/PCAF)^20^. The effects of HAT1 on ISC function display both similarities and differences to the effects of KAT2A/KAT2B loss. *Nguyen* and colleagues created an intestine-specific double KO (DKO) of KAT2A/KAT2 using the tamoxifen-inducible VILLIN-Cre system. Contrary to HAT1^VILLIN KO^ mice, KAT2A/KAT2B DKO mice display a marked decrease in Ki67 and OLFM4 staining in crypts. However, similar to HAT1^VILLIN KO^ mice, KAT2A/KAT2B DKO mice display Lyz2-positive cells throughout villi. In addition, crypt cells from HAT1^VILLIN KO^ mice and KAT2A/KAT2B DKO mice all display significant defects in the formation of intestinal organoids. Interestingly, while loss of KAT2A and KAT2B leads to significant changes in gene expression in the intestine, these histone acetyltransferases may regulate ISC function through multiple mechanisms, including mitochondrial protein acetylation and the regulation of dsRNA accumulation^20^.

Several histone deacetylases (HDACs) are also important for ISC function. Intestinal epithelial cell-specific deletion of HDAC1 and HDAC2 induces increased proliferation of cells in crypts with defects in secretory cell differentiation. Interestingly, HDAC1/2 loss also resulted in striking down-regulation of alpha-defensin gene expression^72^. Examination of the intestines from mice that have a complete deletion of SIRT2 showed similar effects. Loss of SIRT2 led to increased proliferation of crypt cells, an increase in OLFM4 staining, and decreases in secretory cells^17^.

The influence of HAT1 on the structure of LADs is similar to results we had observed in mouse embryonic fibroblasts (MEFs)^67^. In MEFs, HAT1 is required to maintain chromatin accessibility in large domains termed HAT1-dependent accessibility domains (HADs) that overlap with a subset of LADs. This loss of chromatin accessibility is accompanied by an increase in the density of H3K9me3. Our observation that HAT1 primarily effects LADs in intestinal crypt cells suggests that the regulation of LAD structure is a fundamental function of the acetylation of newly synthesized histone H4.

The distribution of LADs throughout the genome of ISCs has not been determined. While a significant fraction of LADs have been found to be conserved across multiple cell types, referred to as constitutive LADs (cLADs), there are also LADs that are cell type-specific (facultative LADs, fLADs)^66,73–75^. Although LADs are typically defined by determining the association of specific sequences with components of the nuclear lamina, LADs are also often distinguished by broad domains of H3K9me3. Our mapping of H3K9me3 in intestinal crypt cells suggests that crypt cells contain fLADs. We detected numerous domains of H3K9me3 in regions of the genome that are not annotated as LADs. Understanding the dynamics of LADs in ISCs, progenitor cells and differentiated cells of the intestine is likely to uncover mechanisms regulating cell type-specific gene expression programs.

Despite significant genome-wide effects on chromatin structure, deletion of HAT1 in the intestine resulted in only modest effects on gene expression. It is unclear whether these changes in gene expression underlie the phenotypic consequences of HAT1 loss in the intestine. It is possible that rather than causing immediate changes in expression, the chromatin changes caused by HAT1 loss influence the ability of cells to activate or deactivate particular genes, biasing differentiation toward particular lineages. HAT1 also has a variety of cellular functions outside of transcriptional regulation that may impact the function and differentiation of ISCs. For example, HAT1 has been shown in multiple organisms to play a role in the repair of DNA double strands breaks and loss of HAT1 results in the accumulation of DNA damage^29,76–80^. DNA damage has been shown to have significant negative impacts on stem cell proliferation and differentiation^81,82^. Loss of HAT1 in MEFs also results in defects in chromosome segregation which may also impair ISC function. Normal mitochondrial function is also necessary for stem cell proliferation and differentiation^83,84^. HAT1 has been shown to be required for the proper mitochondrial function in MEFs and loss of HAT1 results in an increase in ROS and a decrease in the acetylation of several mitochondrial proteins^58,85^. It will be important to decipher the cellular pathways in ISCs that are influenced by HAT1 in future studies.

It is intriguing that HAT1 restricts ISC proliferation *in vivo* while being required for the maintenance of ISC identity and proliferation in organoid cultures *in vitro*. There are several potential explanations for these opposing effects. HAT1 may be integral to signaling pathways that are not precisely replicated in organoid cultures which lack some niche components and microbial metabolites. For example, loss of the BMP receptor *Bmpr1a* increases the proliferation of Lgr5+ ISCs *in vivo* but the loss of BMP signaling *in vitro* disrupts the environment necessary to maintain ISC function in an organoid culture^86^. The proliferative capacity of ISCs may also be distinct from the maintenance of stemness. Deletion of *Mettl3* in the intestine increased the proliferation of ISCs with the formation of elongated crypts. However, Mettl3 loss also impacted the expression of genes involved in the maintenance of stemness, resulting in defects in the formation of organoids^87^. Further studies will be required to unravel the molecular mechanism(s) by which HAT1 regulates ISC proliferation and differentiation.

## Supporting information

Supplemental Marterial

## Acknowledgements

This work was supported by grant R01 GM144601 to MRP and R00 AG05476, DP2CA271361, and Pew Biomedical Sciences Award to MMM.

## Notes

### Competing Interest Statement

The authors have declared no competing interest.

## REFERENCES

1 Beumer, J. & Clevers, H. Cell fate specification and differentiation in the adult mammalian intestine. Nat Rev Mol Cell Biol 22, 39–53 (2021). 10.1038/s41580-020-0278-0

2 Barker, N. Adult intestinal stem cells: critical drivers of epithelial homeostasis and regeneration. Nat Rev Mol Cell Biol 15, 19–33 (2014). 10.1038/nrm3721

3 Clevers, H. The intestinal crypt, a prototype stem cell compartment. Cell 154, 274–284 (2013). 10.1016/j.cell.2013.07.004

4 Jadhav, U. et al. Dynamic Reorganization of Chromatin Accessibility Signatures during Dedifferentiation of Secretory Precursors into Lgr5+ Intestinal Stem Cells. Cell Stem Cell 21, 65–77 e65 (2017). 10.1016/j.stem.2017.05.001

5 Banerjee, K. K. et al. Enhancer, transcriptional, and cell fate plasticity precedes intestinal determination during endoderm development. Genes Dev 32, 1430–1442 (2018). 10.1101/gad.318832.118

6 Kazakevych, J., Sayols, S., Messner, B., Krienke, C. & Soshnikova, N. Dynamic changes in chromatin states during specification and differentiation of adult intestinal stem cells. Nucleic Acids Res 45, 5770–5784 (2017). 10.1093/nar/gkx167

7 Harnik, Y. et al. A spatial expression atlas of the adult human proximal small intestine. Nature 632, 1101–1109 (2024). 10.1038/s41586-024-07793-3

8 Rispal, J., Escaffit, F. & Trouche, D. Chromatin Dynamics in Intestinal Epithelial Homeostasis: A Paradigm of Cell Fate Determination versus Cell Plasticity. Stem Cell Rev Rep 16, 1062–1080 (2020). 10.1007/s12015-020-10055-0

9 Verzi, M. P. & Shivdasani, R. A. Epigenetic regulation of intestinal stem cell differentiation. Am J Physiol Gastrointest Liver Physiol 319, G189–G196 (2020). 10.1152/ajpgi.00084.2020

10 Singh, P. N. P., Madha, S., Leiter, A. B. & Shivdasani, R. A. Cell and chromatin transitions in intestinal stem cell regeneration. Genes Dev 36, 684–698 (2022). 10.1101/gad.349412.122

11 Pivetti, S. et al. Loss of PRC1 activity in different stem cell compartments activates a common transcriptional program with cell type-dependent outcomes. Sci Adv 5, eaav1594 (2019). 10.1126/sciadv.aav1594

12 Chiacchiera, F., Rossi, A., Jammula, S., Zanotti, M. & Pasini, D. PRC2 preserves intestinal progenitors and restricts secretory lineage commitment. EMBO J 35, 2301–2314 (2016). 10.15252/embj.201694550

13 Benoit, Y. D. et al. Polycomb repressive complex 2 impedes intestinal cell terminal differentiation. J Cell Sci 125, 3454–3463 (2012). 10.1242/jcs.102061

14 Koppens, M. A. et al. Deletion of Polycomb Repressive Complex 2 From Mouse Intestine Causes Loss of Stem Cells. Gastroenterology 151, 684–697 e612 (2016). 10.1053/j.gastro.2016.06.020

15 Turgeon, N. et al. The acetylome regulators Hdac1 and Hdac2 differently modulate intestinal epithelial cell dependent homeostatic responses in experimental colitis. Am J Physiol Gastrointest Liver Physiol 306, G594–605 (2014). 10.1152/ajpgi.00393.2013

16 Christopher, H., Zhang, J., Oladejo, S. O., Sharma, S. A. & Kuang, Z. HDAC3 regulates the diurnal rhythms of claudin expression and intestinal permeability. Front Epigenet Epigenom 2 (2024). 10.3389/freae.2024.1496999

17 Li, C. et al. SIRT2 Contributes to the Regulation of Intestinal Cell Proliferation and Differentiation. Cell Mol Gastroenterol Hepatol 10, 43–57 (2020). 10.1016/j.jcmgh.2020.01.004

18 Sheaffer, K. L. et al. DNA methylation is required for the control of stem cell differentiation in the small intestine. Genes Dev 28, 652–664 (2014). 10.1101/gad.230318.113

19 Cedeno, R. J. et al. The histone variant macroH2A confers functional robustness to the intestinal stem cell compartment. PLoS One 12, e0185196 (2017). 10.1371/journal.pone.0185196

20 Nguyen, M. U. et al. KAT2A and KAT2B prevent double-stranded RNA accumulation and interferon signaling to maintain intestinal stem cell renewal. Sci Adv 10, eadl1584 (2024). 10.1126/sciadv.adl1584

21 Takada, Y., Fukuda, A., Chiba, T. & Seno, H. Brg1 plays an essential role in development and homeostasis of the duodenum through regulation of Notch signaling. Development 143, 3532–3539 (2016). 10.1242/dev.141549

22 Holik, A. Z. et al. Brg1 is required for stem cell maintenance in the murine intestinal epithelium in a tissue-specific manner. Stem cells 31, 2457–2466 (2013). 10.1002/stem.1498

23 Liu, S. et al. RPA binds histone H3-H4 and functions in DNA replication-coupled nucleosome assembly. Science 355, 415–420 (2017). 10.1126/science.aah4712

24 Bellelli, R. et al. POLE3-POLE4 Is a Histone H3-H4 Chaperone that Maintains Chromatin Integrity during DNA Replication. Mol Cell 72, 112–126 e115 (2018). 10.1016/j.molcel.2018.08.043

25 Clement, C. & Almouzni, G. MCM2 binding to histones H3-H4 and ASF1 supports a tetramer-to-dimer model for histone inheritance at the replication fork. Nat Struct Mol Biol 22, 587–589 (2015). 10.1038/nsmb.3067

26 Huang, H. et al. A unique binding mode enables MCM2 to chaperone histones H3-H4 at replication forks. Nat Struct Mol Biol 22, 618–626 (2015). 10.1038/nsmb.3055

27 Groth, A. et al. Regulation of replication fork progression through histone supply and demand. Science 318, 1928–1931 (2007). 318/5858/1928 [pii] 10.1126/science.1148992

28 Foltman, M. et al. Eukaryotic replisome components cooperate to process histones during chromosome replication. Cell Rep 3, 892–904 (2013). 10.1016/j.celrep.2013.02.028

29 Nagarajan, P. et al. Histone acetyl transferase 1 is essential for mammalian development, genome stability, and the processing of newly synthesized histones H3 and H4. PLoS Genet 9, e1003518 (2013). 10.1371/journal.pgen.1003518 PGENETICS-D-12-03133 [pii]

30 Parthun, M. R., Widom, J. & Gottschling, D. E. The major cytoplasmic histone acetyltransferase in yeast: links to chromatin replication and histone metabolism. Cell 87, 85–94 (1996). S0092-8674(00)81325-2 [pii]

31 Kleff, S., Andrulis, E. D., Anderson, C. W. & Sternglanz, R. Identification of a gene encoding a yeast histone H4 acetyltransferase. J Biol Chem 270, 24674–24677 (1995).

32 Verreault, A., Kaufman, P. D., Kobayashi, R. & Stillman, B. Nucleosomal DNA regulates the core-histone-binding subunit of the human Hat1 acetyltransferase. Curr Biol 8, 96–108 (1998).

33 Chang, L. et al. Histones in transit: cytosolic histone complexes and diacetylation of H4 during nucleosome assembly in human cells. Biochemistry 36, 469–480 (1997).

34 Schneider, J., Bajwa, P., Johnson, F. C., Bhaumik, S. R. & Shilatifard, A. Rtt109 is required for proper H3K56 acetylation: a chromatin mark associated with the elongating RNA polymerase II. J Biol Chem 281, 37270–37274 (2006).

35 Driscoll, R., Hudson, A. & Jackson, S. P. Yeast Rtt109 promotes genome stability by acetylating histone H3 on lysine 56. Science 315, 649–652 (2007).

36 Han, J. et al. Rtt109 acetylates histone H3 lysine 56 and functions in DNA replication. Science 315, 653–655 (2007).

37 Han, J., Zhou, H., Li, Z., Xu, R. M. & Zhang, Z. Acetylation of lysine 56 of histone H3 catalyzed by RTT109 and regulated by ASF1 is required for replisome integrity. J Biol Chem 282, 28587–28596 (2007).

38 Han, J., Zhou, H., Li, Z., Xu, R. M. & Zhang, Z. The Rtt109-Vps75 histone acetyltransferase complex acetylates non-nucleosomal histone H3. J Biol Chem 282, 14158–14164 (2007).

39 Tsubota, T. et al. Histone H3-K56 acetylation is catalyzed by histone chaperone-dependent complexes. Mol Cell 25, 703–712 (2007).

40 Bazan, J. F. An old HAT in human p300/CBP and yeast Rtt109. Cell Cycle 7, 1884–1886 (2008). 6074 [pii]

41 Berndsen, C. E. et al. Molecular functions of the histone acetyltransferase chaperone complex Rtt109-Vps75. Nat Struct Mol Biol 15, 948–956 (2008).

42 Chen, C. C. et al. Acetylated lysine 56 on histone H3 drives chromatin assembly after repair and signals for the completion of repair. Cell 134, 231–243 (2008). S0092-8674(08)00822-2 [pii] 10.1016/j.cell.2008.06.035

43 Fillingham, J. et al. Chaperone control of the activity and specificity of the histone H3 acetyltransferase Rtt109. Mol Cell Biol 28, 4342–4353 (2008). MCB.00182-08 [pii] 10.1128/MCB.00182-08

44 Das, C., Lucia, M. S., Hansen, K. C. & Tyler, J. K. CBP/p300-mediated acetylation of histone H3 on lysine 56. Nature 459, 113–117 (2009). nature07861 [pii] 10.1038/nature07861

45 Burgess, R. J., Zhou, H., Han, J. & Zhang, Z. A role for Gcn5 in replication-coupled nucleosome assembly. Mol Cell 37, 469–480 (2010). S1097-2765(10)00071-7 [pii] 10.1016/j.molcel.2010.01.020

46 Hoek, M. & Stillman, B. Chromatin assembly factor 1 is essential and couples chromatin assembly to DNA replication in vivo. Proc Natl Acad Sci U S A 100, 12183–12188 (2003).

47 Malay, A. D., Umehara, T., Matsubara-Malay, K., Padmanabhan, B. & Yokoyama, S. Crystal structures of fission yeast histone chaperone Asf1 complexed with the Hip1 B-domain or the Cac2 C terminus. J Biol Chem 283, 14022–14031 (2008). M800594200 [pii] 10.1074/jbc.M800594200

48 Mello, J. A. et al. Human Asf1 and CAF-1 interact and synergize in a repair-coupled nucleosome assembly pathway. EMBO Rep 3, 329–334 (2002).

49 Moggs, J. G. et al. A CAF-1-PCNA-mediated chromatin assembly pathway triggered by sensing DNA damage. Mol Cell Biol 20, 1206–1218. (2000).

50 Ridgway, P. & Almouzni, G. CAF-1 and the inheritance of chromatin states: at the crossroads of DNA replication and repair. J Cell Sci 113, 2647–2658. (2000).

51 Shibahara, K. & Stillman, B. Replication-dependent marking of DNA by PCNA facilitates CAF-1-coupled inheritance of chromatin. Cell 96, 575–585 (1999).

52 Tyler, J. K. et al. Interaction between the Drosophila CAF-1 and ASF1 chromatin assembly factors. Mol Cell Biol 21, 6574–6584 (2001).

53 Liu, W. H., Roemer, S. C., Port, A. M. & Churchill, M. E. CAF-1-induced oligomerization of histones H3/H4 and mutually exclusive interactions with Asf1 guide H3/H4 transitions among histone chaperones and DNA. Nucleic Acids Res 40, 11229–11239 (2012). 10.1093/nar/gks906

54 Winkler, D. D., Zhou, H., Dar, M. A., Zhang, Z. & Luger, K. Yeast CAF-1 assembles histone (H3-H4)2 tetramers prior to DNA deposition. Nucleic Acids Res 40, 10139–10149 (2012). 10.1093/nar/gks812

55 Bhaskara, S. et al. Histone deacetylases 1 and 2 maintain S-phase chromatin and DNA replication fork progression. Epigenetics Chromatin 6, 27 (2013). 10.1186/1756-8935-6-27

56 Sirbu, B. M., Couch, F. B. & Cortez, D. Monitoring the spatiotemporal dynamics of proteins at replication forks and in assembled chromatin using isolation of proteins on nascent DNA. Nat Protoc 7, 594–605 (2012). nprot.2012.010 [pii] 10.1038/nprot.2012.010

57 Zion, E. H. et al. Old and newly synthesized histones are asymmetrically distributed in Drosophila intestinal stem cell divisions. EMBO Rep 24, e56404 (2023). 10.15252/embr.202256404

58 Nagarajan, P. et al. Early-onset aging and mitochondrial defects associated with loss of histone acetyltransferase 1 (Hat1). Aging cell, e12992 (2019). 10.1111/acel.12992

59 Mihaylova, M. M. et al. Fasting Activates Fatty Acid Oxidation to Enhance Intestinal Stem Cell Function during Homeostasis and Aging. Cell Stem Cell 22, 769–778 e764 (2018). 10.1016/j.stem.2018.04.001

60 Langmead, B. & Salzberg, S. L. Fast gapped-read alignment with Bowtie 2. Nat Methods 9, 357–359 (2012). 10.1038/nmeth.1923

61 Wickham, H. ggplot2: Elegant Graphics for Data Analysis. Use R, 1–212 (2009). 10.1007/978-0-387-98141-3

62 Love, M. I., Huber, W. & Anders, S. Moderated estimation of fold change and dispersion for RNA-seq data with DESeq2. Genome Biol 15, 550 (2014). 10.1186/s13059-014-0550-8

63 Illies, C. et al. Requirement of inositol pyrophosphates for full exocytotic capacity in pancreatic beta cells. Science 318, 1299–1302 (2007). 10.1126/science.1146824

64 el Marjou, F., et al. Tissue-specific and inducible Cre-mediated recombination in the gut epithelium. Genesis 39, 186–193 (2004). 10.1002/gene.20042

65 Kaya-Okur, H. S. et al. CUT&Tag for efficient epigenomic profiling of small samples and single cells. Nat Commun 10, 1930 (2019). 10.1038/s41467-019-09982-5

66 Peric-Hupkes, D. et al. Molecular maps of the reorganization of genome-nuclear lamina interactions during differentiation. Mol Cell 38, 603–613 (2010). 10.1016/j.molcel.2010.03.016

67 Popova, L. V. et al. Epigenetic regulation of nuclear lamina-associated heterochromatin by HAT1 and the acetylation of newly synthesized histones. Nucleic Acids Res 49, 12136–12151 (2021). 10.1093/nar/gkab1044

68 Fu, J. et al. Mechanisms and regulation of defensins in host defense. Signal Transduct Target Ther 8, 300 (2023). 10.1038/s41392-023-01553-x

69 Sato, T. et al. Single Lgr5 stem cells build crypt-villus structures in vitro without a mesenchymal niche. Nature 459, 262–265 (2009). 10.1038/nature07935

70 Cheng, C. W., Yilmaz, O. H. & Mihaylova, M. M. Strategies for Measuring Induction of Fatty Acid Oxidation in Intestinal Stem and Progenitor Cells. Methods Mol Biol 2171, 53–64 (2020). 10.1007/978-1-0716-0747-3_4

71 Serra, D. et al. Self-organization and symmetry breaking in intestinal organoid development. Nature 569, 66–72 (2019). 10.1038/s41586-019-1146-y

72 Gonneaud, A. et al. Distinct Roles for Intestinal Epithelial Cell-Specific Hdac1 and Hdac2 in the Regulation of Murine Intestinal Homeostasis. J Cell Physiol 231, 436–448 (2016). 10.1002/jcp.25090

73 Kind, J. et al. Genome-wide maps of nuclear lamina interactions in single human cells. Cell 163, 134–147 (2015). 10.1016/j.cell.2015.08.040

74 Meuleman, W. et al. Constitutive nuclear lamina-genome interactions are highly conserved and associated with A/T-rich sequence. Genome Res 23, 270–280 (2013). 10.1101/gr.141028.112

75 Kind, J. et al. Single-cell dynamics of genome-nuclear lamina interactions. Cell 153, 178–192 (2013). 10.1016/j.cell.2013.02.028

76 Qin, S. & Parthun, M. R. Histone H3 and the histone acetyltransferase Hat1p contribute to DNA double-strand break repair. Mol Cell Biol 22, 8353–8365 (2002).

77 Qin, S. & Parthun, M. R. Recruitment of the type B histone acetyltransferase Hat1p to chromatin is linked to DNA double-strand breaks. Mol Cell Biol 26, 3649–3658 (2006).

78 Yang, X. et al. Histone acetyltransferase 1 promotes homologous recombination in DNA repair by facilitating histone turnover. J Biol Chem 288, 18271–18282 (2013). 10.1074/jbc.M113.473199

79 Tscherner, M., Stappler, E., Hnisz, D. & Kuchler, K. The histone acetyltransferase Hat1 facilitates DNA damage repair and morphogenesis in Candida albicans. Mol Microbiol 86, 1197–1214 (2012). 10.1111/mmi.12051

80 Barman, H. K. et al. Histone acetyltransferase 1 is dispensable for replication-coupled chromatin assembly but contributes to recover DNA damages created following replication blockage in vertebrate cells. Biochem Biophys Res Commun 345, 1547–1557 (2006).

81 Behrens, A., van Deursen, J. M., Rudolph, K. L. & Schumacher, B. Impact of genomic damage and ageing on stem cell function. Nat Cell Biol 16, 201–207 (2014). 10.1038/ncb2928

82 Mani, C., Reddy, P. H. & Palle, K. DNA repair fidelity in stem cell maintenance, health, and disease. Biochim Biophys Acta Mol Basis Dis 1866, 165444 (2020). 10.1016/j.bbadis.2019.03.017

83 Khacho, M. et al. Mitochondrial Dynamics Impacts Stem Cell Identity and Fate Decisions by Regulating a Nuclear Transcriptional Program. Cell Stem Cell 19, 232–247 (2016). 10.1016/j.stem.2016.04.015

84 Wang, Y., Barthez, M. & Chen, D. Mitochondrial regulation in stem cells. Trends Cell Biol 34, 685–694 (2024). 10.1016/j.tcb.2023.10.003

85 Agudelo Garcia, P. A., Nagarajan, P. & Parthun, M. R. Hat1-Dependent Lysine Acetylation Targets Diverse Cellular Functions. J Proteome Res 19, 1663–1673 (2020). 10.1021/acs.jproteome.9b00843

86 Qi, Z. et al. BMP restricts stemness of intestinal Lgr5(+) stem cells by directly suppressing their signature genes. Nat Commun 8, 13824 (2017). 10.1038/ncomms13824

87 Danan, C. H. et al. Intestinal transit-amplifying cells require METTL3 for growth factor signaling and cell survival. JCI Insight 8 (2023). 10.1172/jci.insight.171657

