## Supplemental Marterial for "HAT1 Regulates Intestinal Stem Cell Proliferation and Differentiation"

### SUPPLEMENTARY FIGURE LEGENDS

**Figure S1. Validation of HAT1 Deletion and Phenotypic Analysis.**

**(A)** PCR genotyping of DNA isolated from crypt cells of HAT1$^{villin\ WT}$ and HAT1$^{Villin\ KO}$ mice confirm efficient deletion of the HAT1 gene. **(B)** Western blot analysis of protein lysates from crypt cells of HAT1$^{villin\ WT}$ and HAT1$^{Villin\ KO}$ mice. Blots were probed for HAT1, with Actin as a loading control. **(C)** Quantification of body weight, small intestine length, and colon length comparing WT and KO mice (comparison with student's t-test, N = 5 mice per group).

**Figure S2. HAT1 Deletion Affects Proliferation and Differentiation in the Intestinal Epithelium.**

Representative confocal immunofluorescence images of proximal small intestine sections from HAT1$^{villin\ WT}$ and HAT1$^{Villin\ KO}$ mice following long-term tamoxifen treatment. Sections were stained for Ki67 **(A)**, DCLK1 **(B)**, and Chromogranin A (ChgA **(C)**). Quantification of DCLK1- and ChgA-positive cells per crypt-villus unit is shown (N = 3 mice per group). All images include a 50 µm scale bar.

**Figure S3. H4K12 Acetylation in HAT1-Deficient Intestinal Epithelium.**

Representative confocal immunofluorescence images of proximal small intestine sections from HAT1$^{villin\ WT}$ and HAT1$^{Villin\ KO}$ mice stained for H4K12ac to assess histone H4 lysine 12 acetylation. Images are representative of N = 3 mice per group. Scale bars, 50 µm.

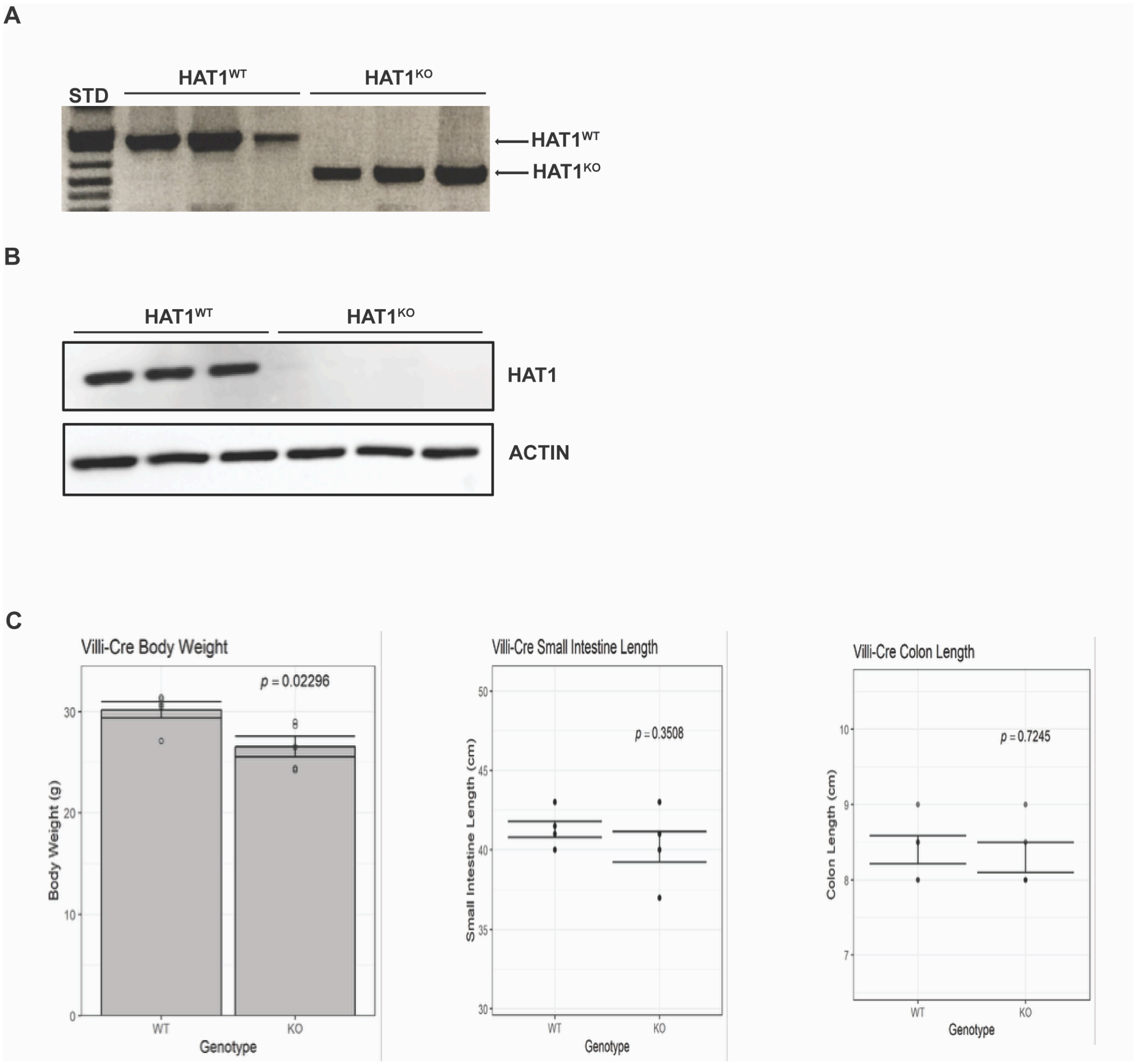
A
STD
HAT1WT
HAT1KO
HAT1WT
HAT1KO
B
HAT1WT
HAT1KO
HAT1
ACTIN
C
Villi-Cre Body Weight
p = 0.02296
Body Weight (g)
WT
KO
Genotype
Villi-Cre Small Intestine Length
p = 0.3508
Small Intestine Length (cm)
WT
KO
Genotype
Villi-Cre Colon Length
p = 0.7245
Colon Length (cm)
WT
KO
Genotype

**Figure S1**

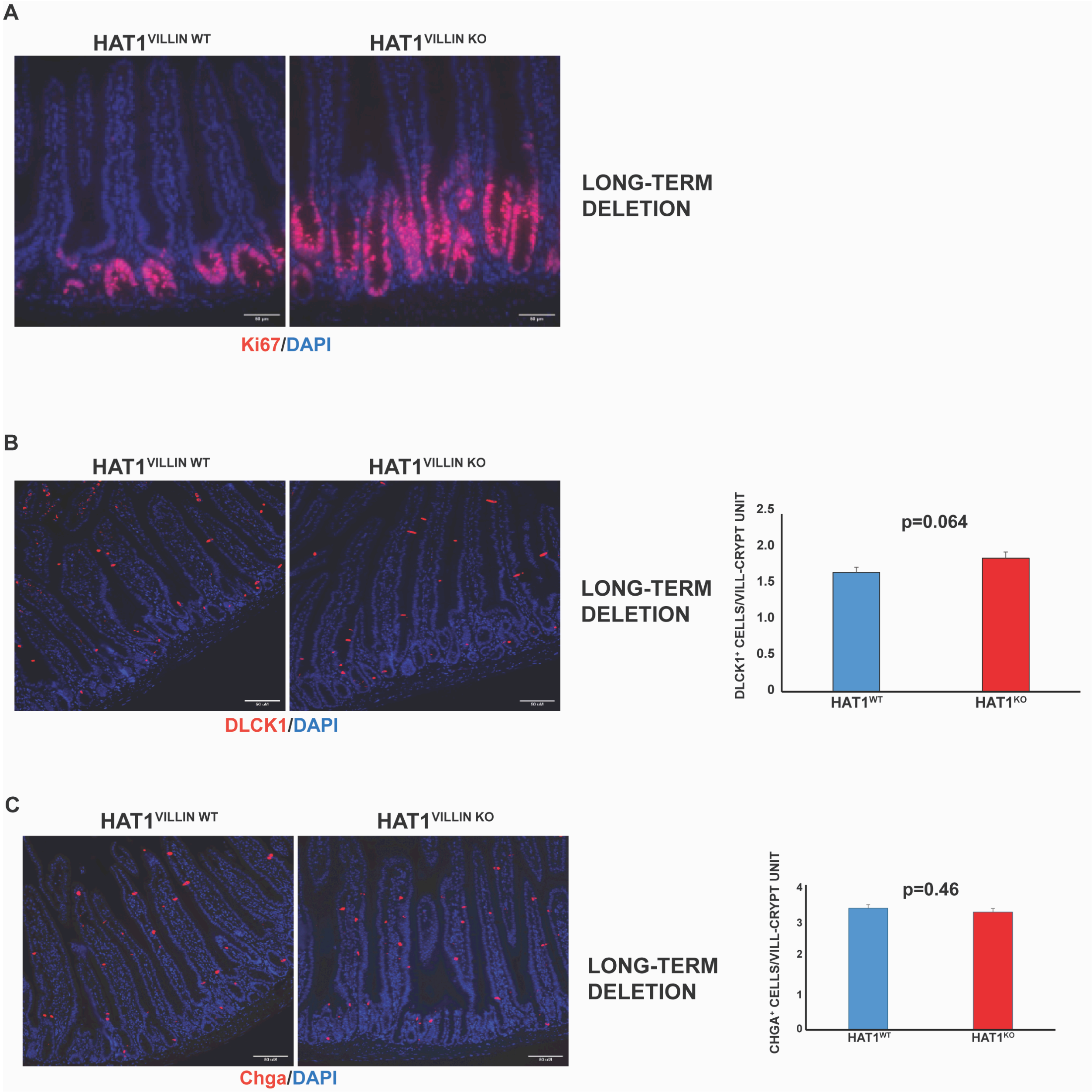
A
HAT1VILLIN WT
HAT1VILLIN KO
LONG-TERM DELETION
Ki67/DAPI
B
HAT1VILLIN WT
HAT1VILLIN KO
LONG-TERM DELETION
DLCK1/DAPI
DLCK1+ CELLS/VILL-CRYPT UNIT
p=0.064
2.5
2.0
1.5
1.0
0.5
0
HAT1WT
HAT1KO
C
HAT1VILLIN WT
HAT1VILLIN KO
LONG-TERM DELETION
Chga/DAPI
CHGA+ CELLS/VILL-CRYPT UNIT
p=0.46
4
3
2
1
0
HAT1WT
HAT1KO

**Figure S2**

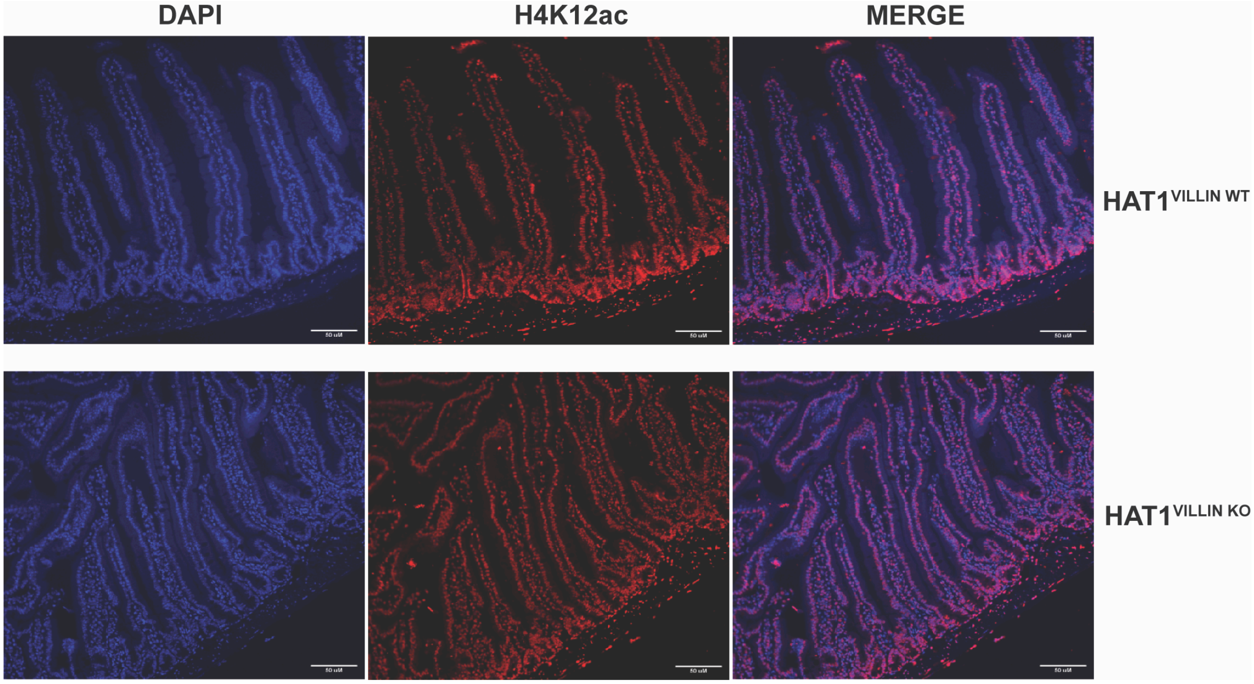


**Figure S3**
